# A Vector Representation of DNA Sequences Using Locality Sensitive Hashing

**DOI:** 10.1101/726729

**Authors:** Lizhen Shi, Bo Chen

## Abstract

Drawing from the analogy between natural language and "genomic sequence language", we explored the applicability of word embeddings in natural language processing (NLP) to represent DNA reads in Metagenomics studies. Here, *k*-mer is the equivalent concept of word in NLP and it has been widely used in analyzing sequence data. However, directly replacing word embedding with *k*-mer embedding is problematic due to two reasons: First, the number of *k*-mers is many times of the number of words in NLP, making the model too big to be useful. Second, sequencing errors create lots of rare *k*-mers (noise), making the model hard to be trained. In this work, we leverage Locality Sensitive Hashing (LSH) to overcoming these challenges. We then adopted the skip-gram with negative sampling model to learn *k*-mer embeddings. Experiments on metagenomic datasets with labels demonstrated that LSH can not only accelerate training time and reduce the memory requirements to store the model, but also achieve higher accuracy than alternative methods. Finally, we demonstrate the trained low-dimensional *k*-mer embeddings can be potentially used for accurate metagenomic read clustering and predict their taxonomy, and this method is robust on reads with high sequencing error rates (12-22%).

## 1 INTRODUCTION

Word embedding is a technique for representing text where different words with similar meaning have a similar real-valued vector representation. It is considered one of the key breakthroughs of machine learning on challenging natural language processing (NLP) problems. There are three popular word embedding models in NLP. Specifically, Global Vectors for words representation (GloVe) [14] from Stanford University uses word-to-word co-occurrence to build the model. word2vec[11] from Google Inc trains a two-layer neural network to reconstruct linguistic contexts of words. FastText [2, 4] is a library developed by the Facebook Research Team for efficient learning of word embeddings and sentence classification. All the three works provide pre-trained vectors for various languages.

The analogy between natural language and "genomic sequence language" has been described earlier [9]. DNA sequences (or reads) are the equivalent concept of sentences and *k*-mers are similar to words in a text document. Metagenomics is the study of a community of microbal species, or the equivalent of a collection of text documents. A metagenome sequencing dataset consists of millions of reads from thousands of species, posing a significant challenge for downstream analyses. NLP techniques may offer a good opportunity to solve problems of this field. However, this is problematic when directly replacing word embedding with *k*-mer embedding due to the following two reasons.

First, lookup tables can be enormous in size due to the large number of *k*-mers. In the word embedding, the lookup table is a embedding weight matrix, which is a two dimensional matrix as shown in figure 1. The space requirement for this matrix is O(*nd*) where *n* is the number of words and *d* is the embedding dimension size. Every row of the embedding matrix is a vector representing a word so every word is represented as a point in the *d* dimensional space. Each word can be converted from a integer to vectors from the embedding matrix where the input integer is the index of a row from the lookup table. The embedding dimension is usually between 100 and 1000 [9]. In a DNA sequence dataset, theoretically there are 4^*k*^ possible *k*-mers, and this number grows exponentially as *k* increases. In practice the number of unique *k*-mer in sequence corpus is much less than the theoretical value, but this number is still far more than the number of words in NLP. Table 1 compares the number of words in English wikipedia and the number of *k*-mers in Pacbio, Sequel, and Nanopore datasets that are used in this paper. The number of *k*-mers in our datasets is 40x the number of words in wikipedia when k=15. Smaller k can not capture useful information. Research suggests that a good accuracy is only achieved with *k*-mers of length at least k = 12 [9](The number of *k*-mers is at least 4^12^=16777216). As a result, applying word embedding models directly to *k*-mers requires hundreds of GB memory and disk to persist a lookup table on a computing node. Such a big table can create significant computational challenges at both training time and test time.

**Figure 1:**
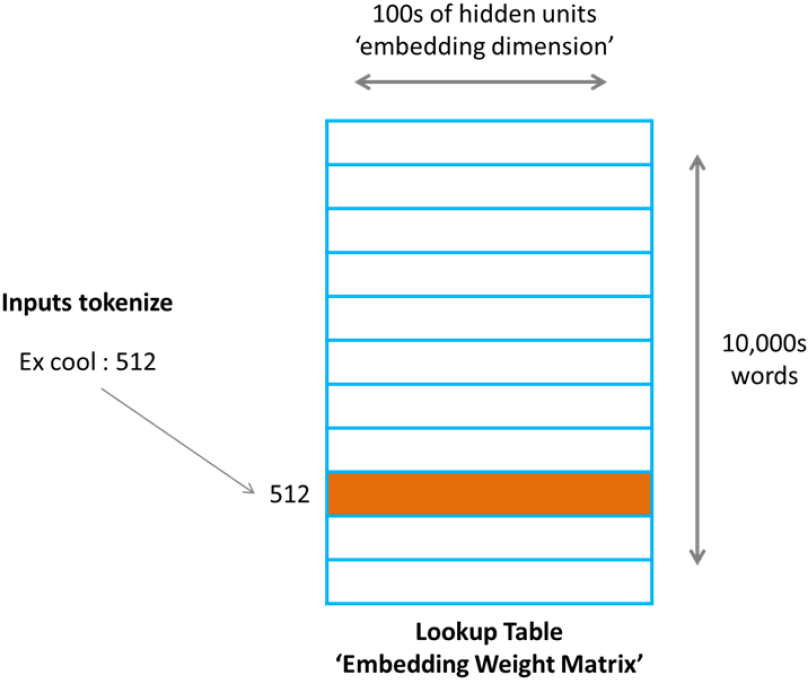
A Lookup Table

**Table 1:** Characteristics of text corpus and DNA sequences

| Data | # Word<br>or $k$-mer* | # of words<br>or $k$-mers* |
| --- | --- | --- |
| Wikipeda-en | 9.7M | 4.6G |
| ActinoMock Pacbio $k=15$ | 468.3M | 2.5G |
| ActinoMock Sequel $k=15$ | 495.6M | 5.3G |
| ActinoMock Nanopore $k=15$ | 436.5M | 3.6G |

\*Calculated by JELLYFISH [8]

Second, the presence of sequencing errors is another challenge. Long-read sequencing technologies like Pacbio and Nanopore sequencing tehcnologies are essential for resolving complex and repetitive regions of the genome in metagenomics. However, they have 10% 15% errors, meaning that the vast majority of the *k*-mers may contain one or two errors for *k* = 15. In word embedding models, the higher frequency of a word in a text corpus, the better vector it can be obtained. Sequencing errors create a lot of rare *k*-mers, which are hard to be trained.

To overcome these challenges, we encode *k*-mers with Locality Sensity Hashing (LSH) in this work. The embedding lookup table size can be efficiently reduced, since LSH can convert large number of *k*-mers to a fixed size of buckets. Those rare *k*-mers generated by sequencing errors can be reduced by the fact that LSH can project similar *k*-mers into the same bucket. After encoding *k*-mers using LSH, we adopted the skip-gram with negative sampling model in NLP to learn *k*-mer embeddings. We further applied our model to solve taxonomic classification problem in metagenomics, where each read must be assigned to a rank in order to obtain a community profile.

To evaluate the quality of our method, we trained embedding and classification models on two metagenomic datasets, which cover a wide range of organisms and taxonomic ranks. We compared the quality of models trained using LSH to those trained on one-hot and FNV encoding using two metrics. First, we test whether we can effectively cluster the reads using their embedding. Second, we tested whether embedding can be used for accurate taxonomic classification at different taxonomic ranks. These experiments demonstrated that our trained embedding vectors are capable of capturing meaningful relationships between reads despite having orders of magnitude fewer dimensions.

In summary, we made the following contributions to the vector representation of DNA sequences:

- We developed a novel method to leverage LSH for encoding genomic sequences. This method can speedup the training time by up to 15.8 times and improves the accuracy by as much as 22.7%, compared to one-hot encoding and FNV encoding.
- We evaluated the impact of read length and error rate on the encoding accuracy.
- We demonstrated that the LSH encoding method can achieve high accuracy in both clustering and classification tasks in metagenomics datasets.

The source code of this work is available at https://github.com/Lizhen0909/LSHVec.

## 2 RELATED WORK

The concept of *k*-mer embedding to represent biological sequences is not a new one. Specifically, bioVec [1] and seq2vec[5] have applied the word2vec technique to biological sequences. Similarly, Gene2vec[3], [21], and Dna2vec[13] applied the same technique to gene embedding, protein embedding, and DNA sequence embedding respectively. All these works were based on word2vec, and their *k*-mer sizes span from 3-8 (dna2vec embed *k*-mers of length 3 to 8, others work on *k*-mer size of 3). The main reason for choosing small *k*-mer is because the *k*-mer size is limited by the lookup table size and computational cost. Large *k*s lead to huge lookup tables and prohibitive computational costs. However, short *k*-mers may not capture high order information such as taxonomy.

Similar to our work, fastDNA[10] was based on *f astT ext* to represent DNA sequence. The *k*-mer size of 8-15 was evaluated in this work. However, it conducted experiments on training database of genomes with error rate less than 5%. Consequently, its performance dropped significantly as the error rate increases[10].

## 3 METHODS

Our implementation is based on *f astT ext*[2, 4], which is an extension and optimization of *word*2*vec*[11]. *f astT ext* is an efficient CPU tool, allowing to train models without requiring a GPU. We made two modifications for DNA encoding: First, we discard the subword feature (aka. *n*-gram) in *f astT ext*, since a (*k − i*)-subword is just a (*k − i*)-mer and an *n*-gram is only a (*k − n* + 1)-mer. This greatly decreases training overhead without losing accuracy. Second, after we obtain *k*-mer embeddings we represent each read by taking the mean of its *k*-mer vectors.

In this following we discuss in details about *k*-mer encoding, *k*-mer embedding (unsupervised), and read classification model (unsupervised).

### 3.1 *k*-mer Encoding

Encoding method in general has a major effect on models’ training time and models’ ability to learn. Our work experimented on 3 encoding methods: One-hot, FNV (Fowler-Noll-Vo), and LSH.

#### 3.1.1 One-hot

A straightforward numeric encoding of *k*-mer is the one-hot encoding, which is widely used in machine learning for turning a categorical feature into a binary vector. Standard one-hot encoding uses 4 bits to encode a nucleotide. To reduce memory overhead, we re-encode each nucleotide in 2 bits (e.g. A,C,G,T to 00, 01, 10 and 11) and then convert the resulting binary vector into an integer. Since most computers use 32 bits to hold an integer, and we only use positive numbers, at most 15-mers can be encoded. A key drawback of one-hot encoding is that the size of lookup table grows exponentially as the k increases, since each *k*-mer corresponds to a specific index number and takes up one row in the lookup table. It is important to note that all the earlier works on the vector representations of biological sequences mentioned implicitly used one-hot as their initial encoding for training their models.

#### 3.1.2 Hash (FNV)

Hashing can be used for converting a large number of *k*-mers into a fixed size of buckets. In this work, we tested FNV (Fowler-Noll-Vo) hashing, a hashing method also used in *f astT ext*.

#### 3.1.3 LSH

Locality Sensitive Hashing (LSH) reduces the dimensionality of high-dimensional data while keeping similar items mapped to the same "buckets" with high probability. LSH has different similarity measures. Choosing the appropriate metric that most suits the problem is very important. We chose the cosine similarity, since it is a good measure of similarity between two vectors in high dimensional space. In our implementation, *A*, *C*, *G*, *T* are encoded as complex numbers of −1, −*i*, *i*, 1. A *k*-mer is represented as a vector in *k*-dimension complex space. Then *n* hyper-planes are drawn from the space and 2^*n*^ buckets are defined. The similarity sim(x,y) of two *k*-mers *x* and *y* is the cosine of the angle between them as shown in the following formula.

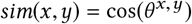

### 3.2 *k*-mer Embedding

We use skip-gram with negative sampling for embedding, a method was originally introduced by [11] and further applied in [2]. Note that the skip-gram model uses the current word to predict its surrounding contextual words, so it is a method of unsupervised learning.

We first constructed a database of *k*-mers and their contexts from our datasets. We defined the ‘context’ as the *k*-mers to the left and to the right of a target *k*-mer within a specific window size. Before training, the embedding matrix was initialized with random weights. The loss function is defined over the entire dataset, and we speedup training with negative sampling, only updating the weight of each target *k*-mer and only a small number (5-20) of negative *k*-mers. Specifically, for a window with *n k*-mers, *w*_*i*_ and *u*_*i*_ are input and output embedding vectors of *k*-mer *i*, training the *k*-mer embedding can be done by minimizing the following loss function:

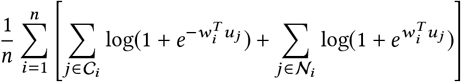

where 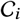 is the indices of *k*-mers in the surrounding context of *k*-mer *w*_*i*_ and 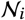 is the indices of negative *k*-mers sampled from the whole *k*-mer set. Please refer to [2] and [11] for more details.

### 3.3 Taxonomy Classification

Taxonomy classification is to assign a given read in a sequence corpus into a fixed number of predefined taxa. Unlike learning of *k*-mer embeddings, taxonomy classification is supervised learning.

Given a set of *n* reads corresponding to *m* labels. Let *w*_*i*_ be the embedding vector of *i*-th *k*-mer, *s*_*j*_ be the embedding vector of *j*-th read. The classification model is a one layer softmax neutral network, trained by minimizing the following objective function:

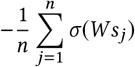

where *σ* is softmax function and *W* is the hidden layer matrix. Refer to [4] for more details.

## 4 DATASETS

We evaluate the quality of our model on two datasets, ActinoMock and CAMI2 Airway (Table 2).

**Table 2:** Statistics of ActinoMock and CAMI2 Airway

| ID | Data | Platform | Insert | Delete | Error | ReadLength |
| --- | --- | --- | --- | --- | --- | --- |
| AM_PB [a] | ActinoMock | Pacbio | 5.4% | 3.7% | 13.0% | 5,947 |
| AM_SEQ [a] | ActinoMock | Sequel | 3.8% | 3.6% | 12.1% | 9,294 |
| AM_NANO [a] | ActinoMock | Nanopore | 2.4% | 3.3% | 12.9% | 17,589 |
| CAMI2_AW [b] | CAMI2_AW | Pacbio | 13.2% | 6.6% | 22.0% | 2,845 |

[a] Calculated by Alfred [15]
[b] From CAMI2 parameters

ActinoMock is a synthetic microbial community using Nanopore, (PacBio) Sequel, and Pacbio (RSII) sequencing platforms. It consists of 12 bacterial strains representing a breadth of genome sizes and ranging from low to high % GC with variable repeat fractions. Taxonomy statistics of this dataset are listed in the supplementary Table 8. We omitted the organism M. coxensis (2623620609) in our analysis due to its negligible number of reads.

The Critical Assessment of Metagenome Interpretation (CAMI) project is the first-ever community-organized benchmark for evaluating computational tools for metagenomes. CAMI is now entering its second-round challenge (CAMI2). Two multisample "toy" data sets are provided to allow participants to prepare for the challenges. We selected 57 organisms from sample 10, 11, and 12 of Human Microbiome Airway dataset and experimented on its Pacbio reads. Table 9 in the supplementary summarized the taxonomy statistics of this dataset.

To avoid imbalanced training data, we subsampled the dataset to select equal number of reads from each species in these two datasets.

## 5 RESULTS

A number of parameters need to be pre-specified in order to train a model. We experimented with different parameters and chosen the followings to train our embedding models: *epoch* = 5, *dim* = 100, and *lr* = 0.05, where *dim* represents embedding dimension, *epoch* is the number of epochs, and *lr* is the learning rate. After learning read embedddings, we projected them onto 2D space for visualization using t-SNE [19].

For classification, we randomly selected 20% of the data as test data and used grid search for parameter tuning. In our experiments it turned out there was little performance difference with *lr* ∈ {1, 0.5}, *epoch* ∈ {15, 20, 25}, *dim* ∈ {50, 100, 200}. We chose a commonly used set of metrics to evaluate multi-class classification [7] performance, including accuracy, precision, recall, f1-score, and support, defined in the following:

- Accuracy is defined as (TP+TN)/(TP+FP+FN+TN) (please refer to Table 3 for TP, FP, FN, and TN).
- Precision is defined as TP/(TP+FP).
- Recall is defined as TP/(TP+FN).
- F1-score is a weighted harmonic mean of the precision and recall, and it is defined as 2*(recall * precision) / (recall + precision)
- Support is the number of occurrences of the true response that lie in each class.

**Table 3:** Evaluation metrics for taxon classification

| Actual Class | Predicted Class |  |  |
| --- | --- | --- | --- |
|  |  | Class = Yes | Class = No |
|  | Class = Yes | True Positive (TP) | False Negative (FN) |
|  | Class = No | False Positive (FP) | True Negative (TN) |

### 5.1 *k*-mer encoding using LSH is more accurate and computational efficient than One-hot and FNV

We compared the accuracy of LSH, FNV, and One-hot encoding by training embedding models on the ActinoMock Nanopore dataset based on these three *k*-mer encoding methods (Methods). The read embedding vectors were projected to 2D space to visualize the separation of different taxa at different ranks (Fig 2). At the Organism level, LSH encoding produces discrete read clusters with each cluster corresponding to a different organism. The only exception is *Halomonas sp. HL-93* (2623620618) and *Halomonas sp. HL-4* (2623620617), the two strains of *Halomonas* that share 99% genomic sequences. Besides that, only very few reads did not get clustered to their genome of origin. The clear separation was similarly observed when the models were trained at the Order, Class and Phylum levels. In contrast, FNV and One-Hot encoding schemes only work well at Phylum level.

**Figure 2:**
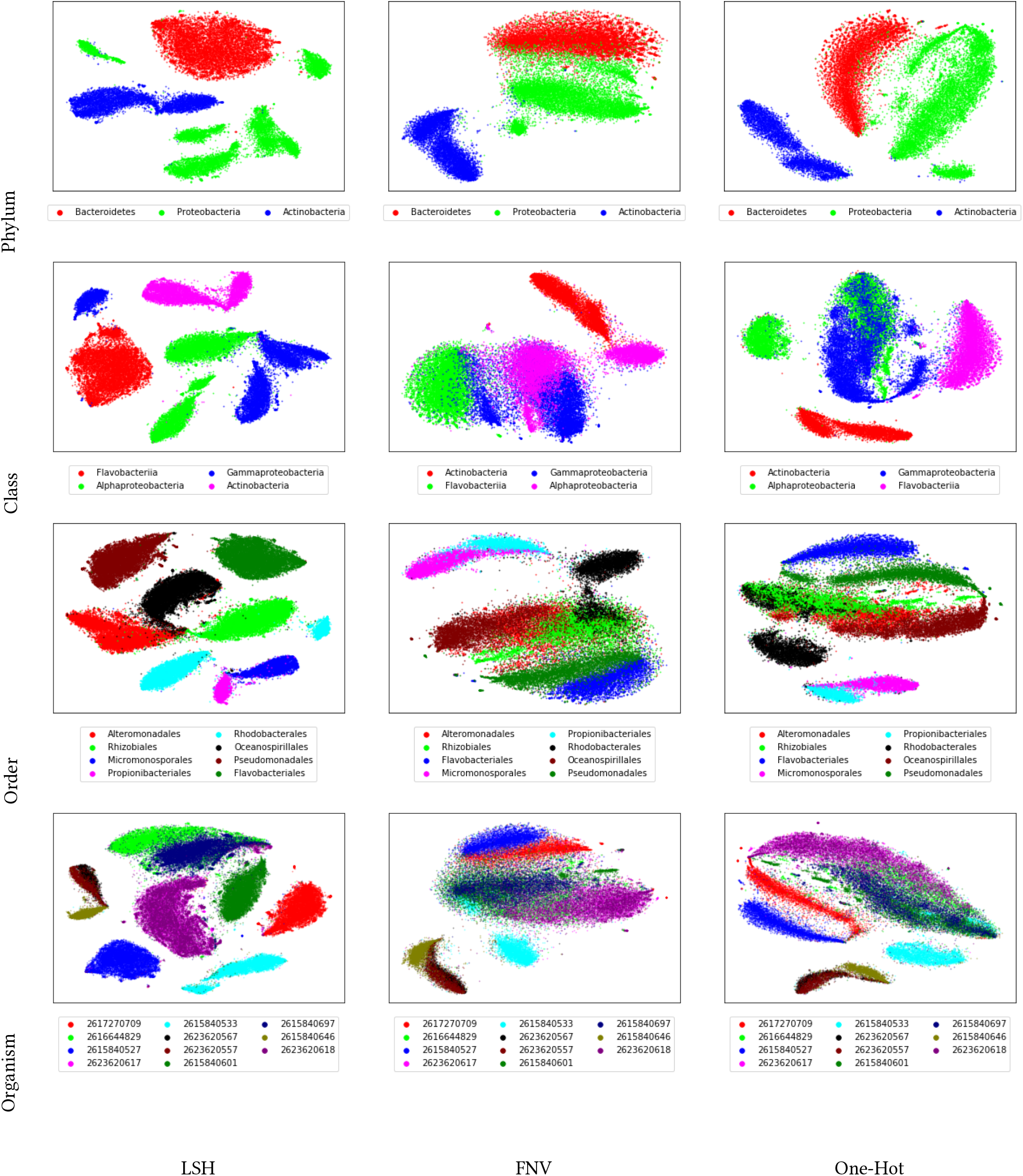
t-SNE visualization for encoding methods

To quantitatively compare the accuracy of these three encoding schemes, we further compared the performance of their taxonomic classification models (Table 4). Consistent with the above clustering visualizations, models trained on LSH outperform the ones trained on one-hot and FNV: better accuracy achieved at all ranks in classification.

**Table 4:**
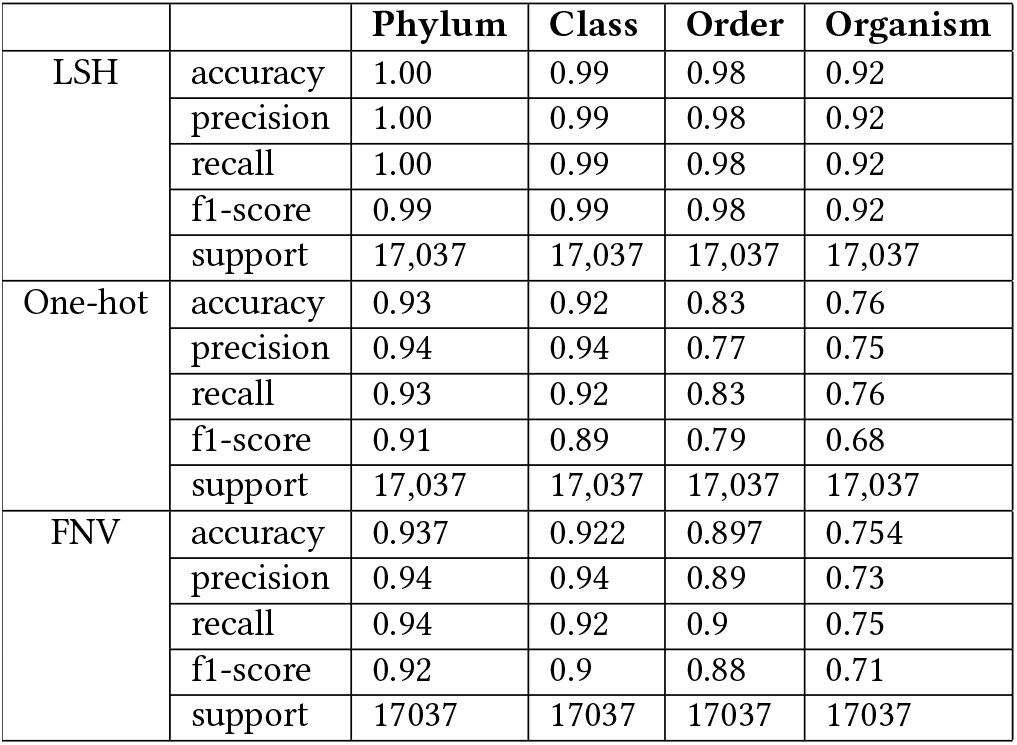
Classification performance of LSH, FNV, and Onehot

|  |  | Phylum | Class | Order | Organism |
| --- | --- | --- | --- | --- | --- |
| LSH | accuracy | 1.00 | 0.99 | 0.98 | 0.92 |
|  | precision | 1.00 | 0.99 | 0.98 | 0.92 |
|  | recall | 1.00 | 0.99 | 0.98 | 0.92 |
|  | f1-score | 0.99 | 0.99 | 0.98 | 0.92 |
|  | support | 17,037 | 17,037 | 17,037 | 17,037 |
| One-hot | accuracy | 0.93 | 0.92 | 0.83 | 0.76 |
|  | precision | 0.94 | 0.94 | 0.77 | 0.75 |
|  | recall | 0.93 | 0.92 | 0.83 | 0.76 |
|  | f1-score | 0.91 | 0.89 | 0.79 | 0.68 |
|  | support | 17,037 | 17,037 | 17,037 | 17,037 |
| FNV | accuracy | 0.937 | 0.922 | 0.897 | 0.754 |
|  | precision | 0.94 | 0.94 | 0.89 | 0.73 |
|  | recall | 0.94 | 0.92 | 0.9 | 0.75 |
|  | f1-score | 0.92 | 0.9 | 0.88 | 0.71 |
|  | support | 17037 | 17037 | 17037 | 17037 |

In theory LSH-encoding and embedding is very memory efficient, as it uses 2 bits to represent one nucleotide instead of 4 bits in onehot. As it only needs one epoch of training while FNV and one-hot need 5, LSH encoding requires much less time. As shown in table 5, the model size of one-hot and FNV (20M) is 15x and 2x respectively of LSH’s, resulting in higher memory consumption. For training speed, LSH takes 40% of One-hot and only 0.6% of FNV’s model training time.

**Table 5:** Computational overhead for different encoding methods

| Hash | ModelSize(Gb) | #words(M) | Mem(Gb) | RT(hr) |
| --- | --- | --- | --- | --- |
| FNV 20M | 20 | 20 | 11 | 19 |
| One-hot | 130 | 360 | 180 | 3 |
| LSH 25bit | 8.5 | 9 | 6 | 1.2 |

### 5.2 Our approach is able to work well with different sequencing technologies

Existing sequencing technologies display platform-specific biases depending on run mode and chemistry. These biases affect read length, data throughput, GC coverage bias, the ability to resolve repetitive genomic elements and error rates [6, 12, 16]. In order to evaluate the applicability of our methods to these technologies, we trained models on Pacbio, Sequel, and Nanopore reads respectively of ActinoMock dataset using our proposed LSH encoding. Figure 3 presented the visualization of our trained embedding models on the order rank, where clear clusters can be observed. We didn’t show the result on the organism rank, because there exist some overlaps: some organisms(2623620617 and 2623620618, 2623620557 and 2623620567, and 2616644829 and 2615840697) share a great many DNA sequences and are from the same species as shown in the supplementary table 8. Table 6 shows the classification accuracy on the order rank. These results suggest that the reads were assigned accurately with the trained classification models.

**Figure 3:**
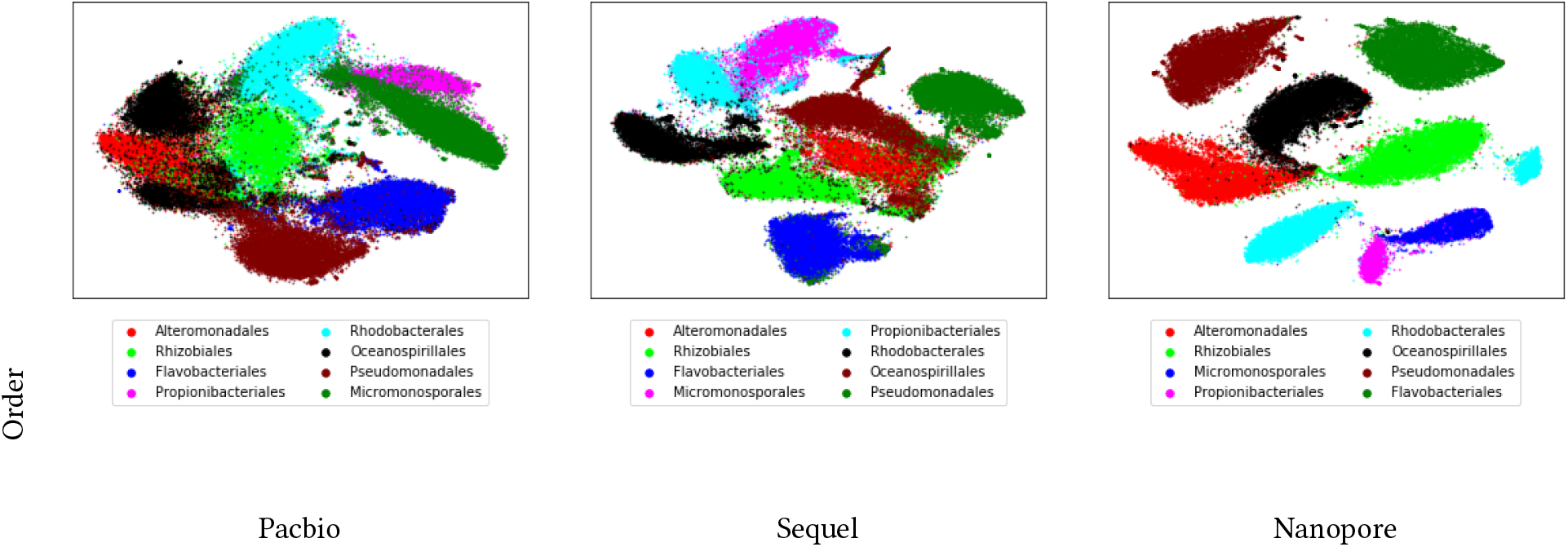
t-SNE visualization of Pacbio, Sequel, and Nanopore reads on ActinoMock dataset

**Table 6:** Classification performance of Pacbio, Sequel, and Nanopore reads on the order rank

|  | accuracy | precision | recall | f1-score | support |
| --- | --- | --- | --- | --- | --- |
| Nanopore | 0.98 | 0.98 | 0.98 | 0.98 | 17,037 |
| Sequel | 0.92 | 0.92 | 0.92 | 0.92 | 20,935 |
| Pacbio | 0.88 | 0.88 | 0.88 | 0.88 | 19,919 |

In terms of these sequencing technologies, we can see that Nanopore achieved the best result, while Pacbio achieved the worst. This is likely caused by the fact that Nanopore has the longest read length and lowest error rate (as shown in Table 2). The impact of read length and error rate on the model’s accuracy was evaluated in the next part.

### 5.3 The impact of read length and error rate on the model’s accuracy

Read length and error rate are two important characteristics of DNA reads and hence can have major impact on the model performance. Intuitively, longer length and lower error rate enable more accurate estimation and lead to higher model accuracy, since longer reads convey more information, reads with lower error rates are more accurate. In order to evaluate the impact of read length and error rate on the model’s accuracy, we simulated several datasets with various number of read lengths and error rates from the synthetic ActinoMock metagenome dataset using the CAMI2 Pacbio simulator. Figure 4 showed the result of our experiments. As expected, the model accuracy was falling as the error rate increased and the read length decreased.

**Figure 4:**
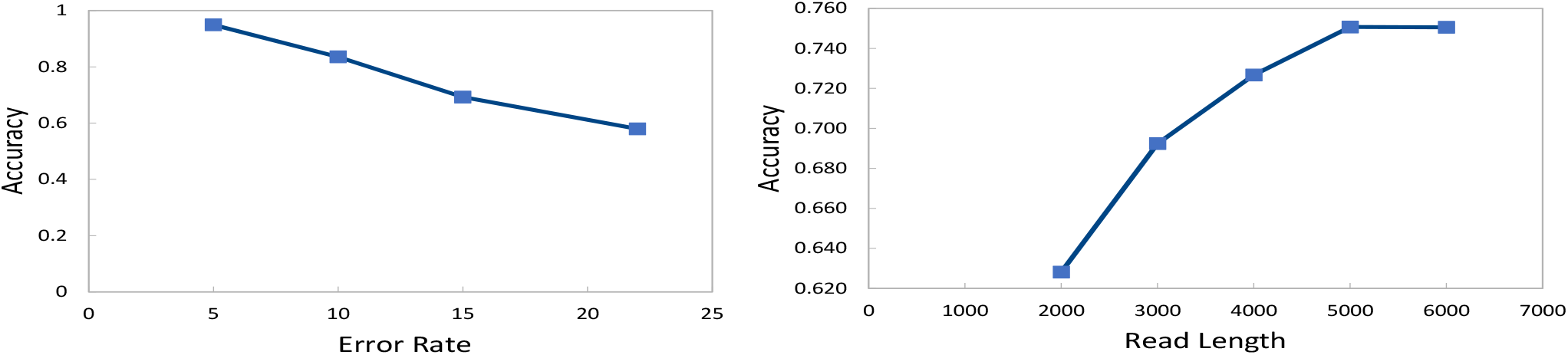
The impact of read length and error rate on model accuracy

### 5.4 Our approach works well on the complex dataset

Clustering and classification on complex community metagenomic data remains a challenging issue for environmental studies and usually achieves very low accuracy. In order to evaluate the efficacy of our methods on complex dataset, we ran it on CAMI2 Airway. The visualization of the embedding models was presented in Fig 5. Two experiments were conducted for classification: First, we randomly sampled 20% of reads as test data (CAMI2_AW); Second, we sampled 20% of organisms, choosing 12 out of 57 organisms (starred in Table 9) as test data (*CAMI2_AW), since in reality, we either label or not label all the reads belonging to an organism dependent on whether the organism is known or not. The classification accuracy is summarized in table 7. The performance of *CAMI2_AW is 1% to 5% worse than that of CAMI2_AW in f1-score. The most probable reason is that randomly splitting high-coverage reads may introduce overfitting.

**Figure 5:**
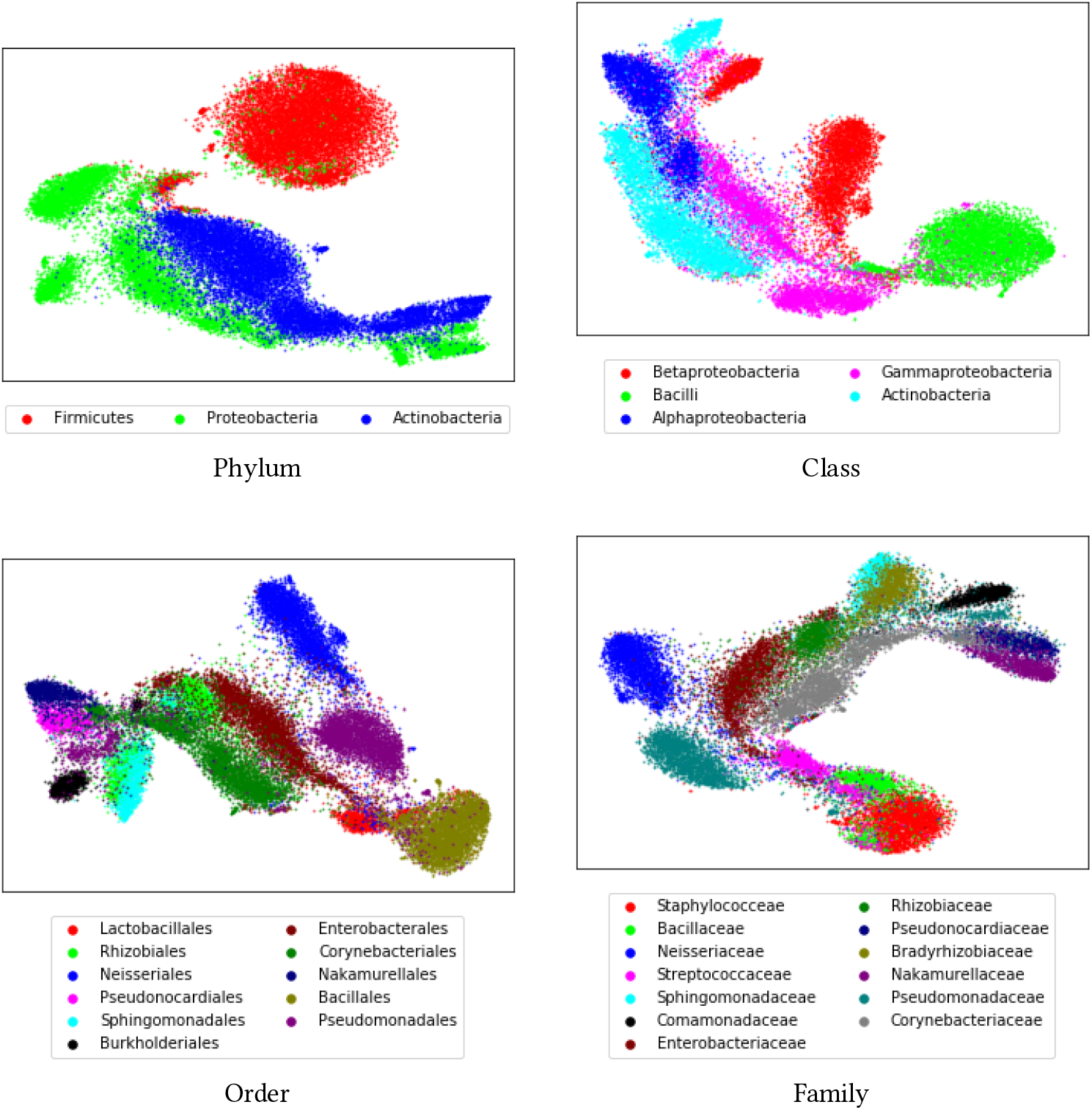
t-SNE visualization for CAMI2 Airway dataset

**Table 7:** Classfication tests for CAMI2 Airway dataset

|  |  | Phylum | Class | Order | Family |
| --- | --- | --- | --- | --- | --- |
| 5*CAMI2_AW | accuracy | 0.92 | 0.90 | 0.86 | 0.85 |
|  | precision | 0.92 | 0.89 | 0.86 | 0.85 |
|  | recall | 0.92 | 0.90 | 0.86 | 0.85 |
|  | f1-score | 0.92 | 0.90 | 0.86 | 0.85 |
|  | support | 120,000 | 120,000 | 120,000 | 120,000 |
| 5**CAMI2_AW | accuracy | 0.87 | 0.83 | 0.72 | 0.72 |
|  | precision | 0.87 | 0.85 | 0.82 | 0.84 |
|  | recall | 0.87 | 0.83 | 0.87 | 0.86 |
|  | f1-score | 0.87 | 0.83 | 0.84 | 0.84 |
|  | support | 120,000 | 120,000 | 100,000 | 100,000 |

Overall, the results on CAMI2 Airway exhibits lower accuracy than that on ActinoMock, this is likely caused by two reasons: 1) The reads of CAMI2 Airway are shorter and has more errors (as shown in Table 2); 2) CAMI2 Airway dataset is much more complex than ActinoMock dataset (ActinoMock has 12 organisms while CAMI2 airway has 57).

## 6 DISCUSSION AND CONCLUSION

In this work, we presented a simple but effective vector representation model by using locality sensitive hashing and skip-gram with negative sampling. LSH significantly reduces the lookup table size and enhances training power by keeping similar *k*-mers in one bucket. Comparing LSH to one-hot and FNV proves LSH has advantages in terms of accuracy and resource costs.

To evaluate the quality of our method, we trained embedding models and classification models on both real mock and simulating metagenomic dataset: ActinoMock and CAMI2 Airway. The trained read vectors verified the promising performance of our method to capture the rich characteristics of DNA sequences despite having orders of magnitude fewer dimensions.

Graph has a longstanding place in biological sequence analysis, either de bruijn graph [20] or read graph [17]. Common graph techniques even with big data technologies [18] do not scale well when the graph size increases to a certain degree, which prevents biological scientist from gaining insights into biological sequences. With the reads embedding, we opened a new door to genome analysis. Various numerical-based techniques can be applied, which are more mature and able to scale better than graph. In particular, the learned vectors through reads embeddings can be fed to various machine learning models for applications in bioinformatics.

However, taxonomic classification belongs to the category of supervised learning, which means that it is reliant on taxonomy reference databases for classifying sequences. Despite the ever growing sequence databases, most metagenomic reads cannot be assigned to a function, limiting the value of metagenomic datasets as a tool for novel discoveries.

It is important to note that we demonstrated our method with genome analysis in this work. However, We believe our method is also applicable to other biological sequences, like protein embedding and gene embedding. We opened source our implementation to facilitate comparison of future work and boost the application of word embedding in Bioinformatics.

## ACKNOWLEDGMENTS

Grid search of parameters for this project was performed on the HPC cluster at the Research Computing Center at the Florida State University (FSU).

## A SUPPLEMENTARY MATERIAL

**Table 8:** Reference statistics of ActionoMock obtaining from IMG database

| IMG Tax | Organism | Phylum | Class | Order | Family | Genus |
| --- | --- | --- | --- | --- | --- | --- |
| 2623620557 | Micromonospora echinaurantiaca DSM 43904 | Actinobacteria | Actinobacteria | Micromonosporales | Micromonosporaceae | Micromonospora |
| 2623620567 | Micromonospora echinofusca DSM 43913 | Actinobacteria | Actinobacteria | Micromonosporales | Micromonosporaceae | Micromonospora |
| 2615840646 | Propionibacteriaceae bacterium ES.041 | Actinobacteria | Actinobacteria | Propionibacteriales | Propionibacteriaceae |  |
| 2615840527 | Muricauda sp. ES.050 | Bacteroidetes | Flavobacteriia | Flavobacteriales | Flavobacteriaceae | Muricauda |
| 2615840601 | Cohaesibacter sp. ES.047 | Proteobacteria | Alphaproteobacteria | Rhizobiales | Cohaesibacteraceae | Cohaesibacter |
| 2615840533 | Thioclava sp. ES.032 | Proteobacteria | Alphaproteobacteria | Rhodobacterales | Rhodobacteraceae | Thioclava |
| 2623620617 | Halomonas sp. HL-4 | Proteobacteria | Gammaproteobacteria | Oceanospirillales | Halomonadaceae | Halomonas |
| 2623620618 | Halomonas sp. HL-93 | Proteobacteria | Gammaproteobacteria | Oceanospirillales | Halomonadaceae | Halomonas |
| 2616644829 | Marinobacter sp. LV10MA510-1 | Proteobacteria | Gammaproteobacteria | Alteromonadales | Alteromonadaceae | Marinobacter |
| 2615840697 | Marinobacter sp. LV10R510-8 | Proteobacteria | Gammaproteobacteria | Alteromonadales | Alteromonadaceae | Marinobacter |
| 2617270709 | Psychrobacter sp. LV10R520-6 | Proteobacteria | Gammaproteobacteria | Pseudomonadales | Moraxellaceae | Psychrobacter |

**Table 9:**
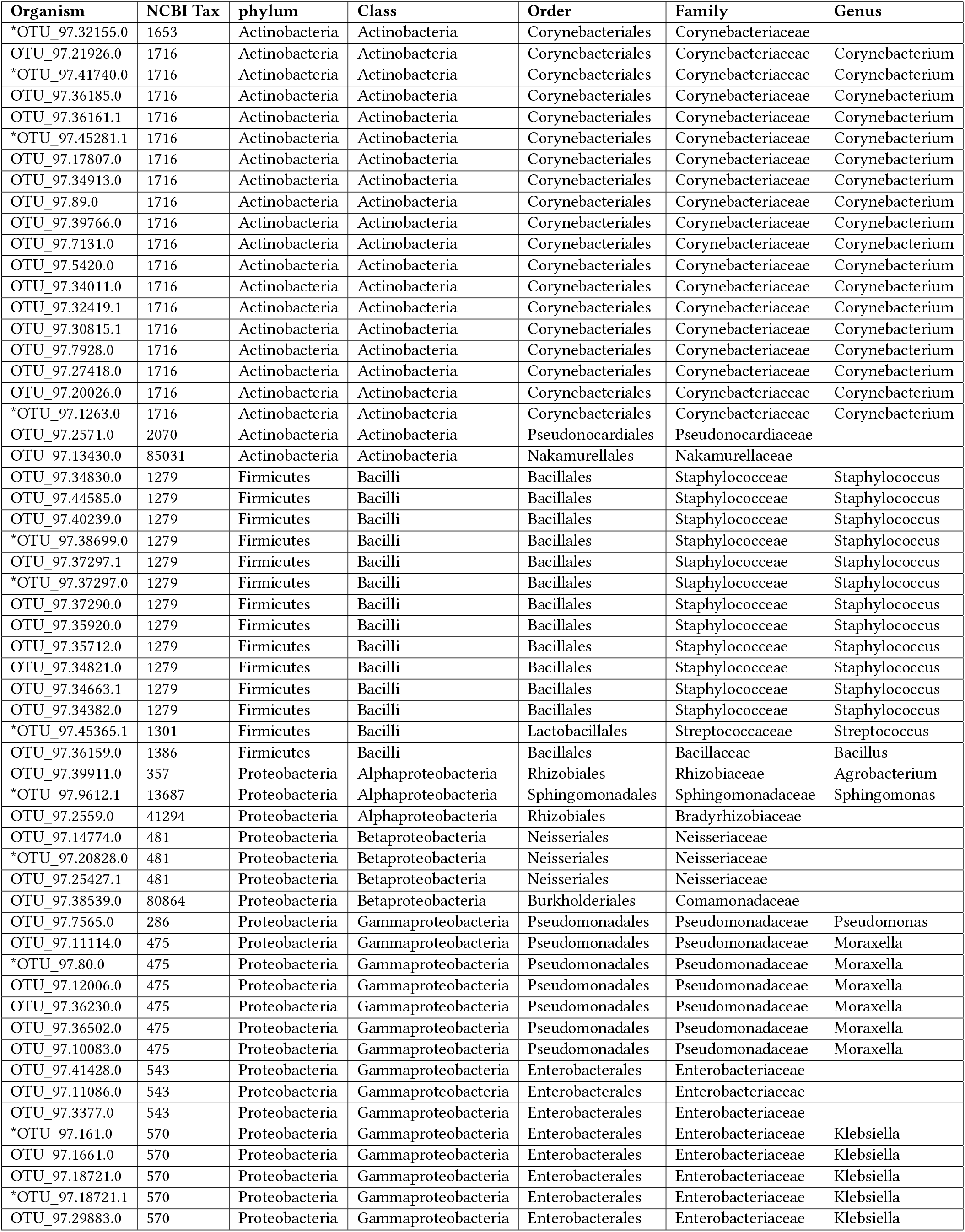
Taxonomy Statistics of CAMI2 AirWay

